# Gaze direction as equilibrium: more evidence from spatial and temporal aspects of small-saccade triggering in the rhesus macaque monkey

**DOI:** 10.1101/766212

**Authors:** Ziad M. Hafed, Laurent Goffart

## Abstract

Rigorous behavioral studies made in human subjects have shown that small-eccentricity target displacements are associated with increased saccadic reaction times, but the reasons for this remain unclear. Before characterizing the neurophysiological foundations underlying this relationship between the spatial and temporal aspects of saccades, we tested the triggering of small saccades in the male rhesus macaque monkey. We also compared our results to those obtained in human subjects, both from the existing literature and through our own additional measurements. Using a variety of behavioral tasks exercising visual and non-visual guidance of small saccades, we found that small saccades consistently require more time than larger saccades to be triggered in the non-human primate, even in the absence of any visual guidance and when valid advance information about the saccade landing position is available. We also found a strong asymmetry in the reaction times of small upward versus downward visually-guided saccades, similar to larger saccades, a phenomenon that has not been described before for small saccades, even in humans. Following the suggestion that an eye movement is not initiated as long as the visuo-oculomotor system is within a state of balance, in which opposing commands counterbalance each other, we propose that the longer reaction times are a signature of enhanced times needed to create the symmetry-breaking condition that puts downstream premotor neurons into a push-pull regime necessary for rotating the eyeballs. Our results provide an important catalog of non-human primate oculomotor capabilities on the miniature scale, allowing concrete predictions on underlying neurophysiological mechanisms.

## Introduction

The sudden appearance of a visual target is most of the time followed by a saccadic movement of the eyes. In non-pathological conditions, this movement brings the image of the target within the central visual field. During the subsequent fixation, small saccades can still be triggered, even though the target location in space has not changed. In the majority of studies using monkeys as behavioral research subjects, so-called computer-controlled “fixation windows” are used to make sure that the animal effectively looks at the fixated target and not elsewhere. Such windows can constrain the range of saccade sizes that the monkey is allowed to make during fixation. If the extent of the window is very small, then the propensity to generate saccades bringing gaze direction outside its boundaries will inevitably be reduced. However, the generation of fixational saccades in the monkey is not a mere function of computer-controlled constraints on fixation accuracy. Their amplitude remains small even when large fixation windows are used (e.g. Guerrasio et al. 2010). Moreover, high acuity visual tasks often require that small saccades are directed in highly precise manners, and monkeys can make microsaccades that accurately and consistently orient a restricted zone of their retina toward the location of tiny visual spots (Tian et al. 2018; 2016). Another aspect that influences the generation of “fixational” saccades is the target size. Minuscule targets indeed elicit saccades whose range of amplitudes is smaller than larger targets (Goffart et al. 2012).

Besides these spatial aspects, there are also temporal aspects, such as variabilities in the timing of saccade generation. From the excitation of ganglion cells in the retina to the recruitment of motor neurons and the contraction of extraocular muscles, action potentials are transmitted through several relays in the cerebrum (thalamus, cerebral cortex, superior colliculus, and reticular formation). The latency of saccades reflects the time (duration) taken by the action potentials to recruit a sufficient number of neurons to contract the agonist muscles while relaxing the antagonist ones, and rotate the eyeballs. Thus, any lesion that compromises the visuomotor transmission leads to increasing the oculomotor reaction time. All other things being kept equal, the visuomotor delay in humans depends upon the eccentricity of the target in the visual field (Kalesnykas and Hallett 1996; 1994; Wyman and Steinman 1973) insofar as the latency of saccades towards foveal targets is much longer than other saccades. However, since these observations were made, it was not entirely clear whether the origin of these longer latencies was visual or motor in origin. Later experiments testing saccades towards auditory targets (Zambarbieri et al. 1995) or gaze shifts that were rendered dysmetric by a cerebellar pharmacological perturbation (Goffart and Pelisson 1997) suggested that the dependency was motor-related: the smaller the saccade, the longer the time to initiate it. We hypothesize that this effect is related to a recent proposal that gaze direction is an equilibrium, and that an eye movement (saccadic or slow) is not initiated as long as the visuo-oculomotor system is within a mode where opposing commands counterbalance each other (Goffart 2019; Goffart et al. 2018; Krauzlis et al. 2017).

Here we document the timing of saccade triggering in rhesus macaque monkeys in a variety of behavioral tasks. We particularly focus on very small saccades, as well as differences between different saccade directions, in order to investigate hypotheses related to recent neurophysiological findings (Chen et al. 2019; Guerrasio et al. 2010; Hafed and Chen 2016; Krauzlis et al. 2017) and also motivate future ones. Our results are consistent with the model positing that saccade triggering depends on balance of different opposing oculomotor commands, and with particular dependence on spatial visuomotor maps magnifying the representation of the central visual field. Our analyses of small-saccade triggering in this paper provide an important documentation of oculomotor capabilities of a popular animal model used for neurophysiological investigations of the primate oculomotor system, and they provide constraints on interpreting the existing neurophysiological literature as well as hypotheses for further mechanistic studies.

## Methods

### Ethics approvals

All monkey experiments were approved by ethics committees at the Regierungspräsidium Tübingen. The experiments were in line with the European Union directives and the German laws governing animal research. Some monkey data were analyzed from (Willeke et al. 2019) for the new purposes of this article. In these cases, the same committees had approved the experiments.

We also analyzed anew human data from the same study (Willeke et al. 2019), as well as collected additional data from one author (ZH). These human experiments were approved by ethics committees at the Medical Faculty of Tübingen University.

### Laboratory setups

Monkey experiments were performed in the same laboratory environment as that described recently (Buonocore et al. 2019; Chen and Hafed 2018; Chen et al. 2018; Skinner et al. 2019; Willeke et al. 2019). Human experiments were done in the laboratory described in (Grujic et al. 2018; Hafed 2013).

Monkey eye movements were recorded at 1kHz using electromagnetic induction (Fuchs and Robinson 1966; Judge et al. 1980). As stated in (Willeke et al. 2019), we used video-based eye tracking for the human subjects.

### Animal preparation

We collected behavioral data from 2 adult, male rhesus macaques (Macaca Mulatta). Monkeys M and N (aged 7 and 10 years, and weighing 8 and 11.5 kg, respectively) were implanted with a scleral search coil to allow measuring eye movements using the electromagnetic induction technique (Fuchs and Robinson 1966; Judge et al. 1980). The monkeys were also implanted with a head holder to stabilize their head during the experiments, with details on all implant surgeries provided earlier (Chen and Hafed 2013; Skinner et al. 2019). They were part of a larger neurophysiology project beyond the scope of the current manuscript.

### Monkey behavioral tasks

The monkeys were trained to perform a visually-guided saccade task. Each trial started with the presentation of a central white fixation spot (86 cd/m^2^) over a uniform gray background (29.7 cd/m^2^). The fixation spot was a square of 5.3 x 5.3 min arc dimensions. After 300-900 ms of fixation (i.e. maintaining eye position within a prescribed distance from the spot; see below), the fixation spot was jumped to a new location, instructing the monkeys to generate a visually-guided saccade to follow the spot. The size of the jump was varied randomly across trials. Target locations were chosen from among 96 predefined possibilities, as follows: the target could jump by a distance of 0.06, 0.1, 0.2, 0.3, 0.5, 0.7, 1, 1.5, 2, 3, 5, or 10 deg in either the horizontal or vertical dimension, or both simultaneously. Moreover, the jump could be positive (i.e. rightward or upward) or negative (i.e. leftward or downward) from display center. Therefore, we sampled horizontal, vertical, and diagonal target locations of different eccentricities, with denser sampling for foveal locations. If the monkeys fixated the new spot location within 500 ms after it had jumped, and held their eye position there for another approximately 300 ms, they were rewarded with liquid reward.

We controlled the monkeys’ fluid reward system in real-time by employing a virtual, computer-controlled window around target location. If eye position entered the virtual window within the prescribed “grace” period, a reward was triggered. Otherwise, the trial was aborted, and a new trial was initiated. Our virtual “target windows” across trials had radii of 2-2.5 deg. Note that a radius of 2-2.5 deg was still employed even for foveal target locations of smaller eccentricities. This means that for such small target eccentricities, we exploited the natural tendency of the monkeys to perform the task without any computer monitoring to ensure that they generated the required saccades. This was not a problem at all, because after the monkeys were trained on the task with eccentricities of 5 deg and higher, they very naturally generalized their trained rule when tested on smaller target eccentricities. This was also the case in more complicated variants of the task (Willeke et al. 2019), and it was also consistent with human results (e.g. see Fig. 9). We felt that this approach of large virtual target windows was better than the alternative of enforcing tiny target windows, because in the latter case, any potential increases in reaction times of saccades could have been interpreted as being the consequence of increased task difficulty.

We analyzed a total of 928 trials from monkey M in this task, and 1246 trials from monkey N.

We also analyzed delayed, visually-guided saccades and memory-guided saccades made by the same two monkeys. These data were collected during an earlier experiment, with detailed methods described elsewhere (Willeke et al. 2019). The purpose of the present re-analysis was to explore saccade latency as a function of target eccentricity, and to examine how this relationship might be affected by task instruction. We also wanted to directly compare results from the same animals used in the (immediate) visually-guided saccade experiments described above. Briefly, the delayed saccade task was similar to the (immediate) visually-guided saccade task described above, except that there was a time interval during which the fixation spot remained visible when the saccade target was visible. The presence of the central spot instructed the monkeys to maintain fixation, despite the presence of the peripheral target. When the fixation spot disappeared, the monkeys could make the saccade to the peripheral target. This task allowed us to investigate whether increased saccadic latencies for small target eccentricities (see Results) were necessarily linked to sudden visual onsets in the (immediate) visually-guided saccade task.

The memory-guided saccade task was similar to the delayed, visually-guided saccade task, except that the target duration was brief (duration: 58 ms). Thus, practically the entire delay period (300-1100 ms after the target flash) had only the fixation spot visible. When the fixation spot disappeared, the monkeys generated an eye movement to the remembered location of the earlier target flash. This task was useful to dissociate increased reaction times for small target eccentricities from the presence of a visual target.

In all tasks, we started out with the monkeys already being experts in oculomotor tasks requiring fixation of a small target (similar to author ZH in Fig. 9). The monkeys were used in earlier studies demonstrating their level of precision in eye movement control (e.g. Tian et al. 2018; 2016 for monkey N and Buonocore et al. 2019; Skinner et al. 2019 for monkey M). Therefore, even though we analyzed thousands of trials in the present study, we could not characterize learning processes. What we can say, however, is that when the monkeys were completely naïve, we first showed them similarly sized small fixation spots and they naturally fixated them to a precision significantly higher than that required by computer-controlled virtual windows. The quality of their fixation therefore started out being good, and improved fairly quickly within a matter of a few trials within a single session.

### Human behavioral tasks

For supporting comparisons of the results from our monkeys in the (immediate) visually-guided saccade task to those reported in the literature on human subjects (e.g. Kalesnykas and Hallett 1996; 1994; Wyman and Steinman 1973), we ran one human subject (author ZH) on the same task as that performed by our two expert monkeys (see Fig. 9). We analyzed 1966 trials from this subject. In separate sessions, we then ran a variant of the same task, but the fixation spot now remained visible after target jump. The subjects’ task was to maintain fixation and press a button (with the right thumb) as quickly as possible after target onset. The goal was to measure manual reaction times for perceptual detections not requiring an eye movement. This allowed us to compare and contrast manual reaction times to saccadic reaction times from the original variant of the task. We analyzed 1924 from the same subject in this task variant.

Because we were particularly interested in the monkey memory-guided saccade reaction time results as a function of target eccentricity (see Results below), we decided to explore their generalizability to human memory-guided saccades, an aspect that was not well-explored in the existing human saccade literature so far. Therefore, we re-analyzed human data that we had collected earlier (Willeke et al. 2019) with the same task. Briefly, the human subjects made the same memory-guided saccade task with target locations being chosen randomly across trials from among 480 possibilities, with heavy sampling of small eccentricities.

### Behavioral analyses

We detected saccades and microsaccades using established methods reported elsewhere (Bellet et al. 2019; Chen and Hafed 2013). We manually inspected each trial to correct for false alarms or misses by the automatic algorithms, which were rare. We also marked blinks or noise artifacts for later removal.

In the (immediate) visually-guided saccade task, we analyzed the saccade that was triggered after the target jump. We excluded all trials in which there was a blink within +/- 100 ms from target jump, since this could impair visual detection of the jump. We also excluded all trials in which a microsaccade occurred within the period from −100 ms to 60 ms relative to target jump occurrence. Our reason was that microsaccades around stimulus onset reduces target visibility and increases reaction time (Beeler 1967; Bellet et al. 2017; Chen and Hafed 2017; Chen et al. 2015; Hafed and Krauzlis 2010; Tian et al. 2016; Zuber and Stark 1966). We defined as a successful reaction any eye movement made within 70-500 ms after target jump (throughout this article, we interchangeably refer to the time interval between target jump and saccade onset as the “saccadic latency” or the “saccadic reaction time”). When plotting reaction time as a function of target eccentricity or direction, or both, we binned nearby eccentricities appropriately, and we only showed summary measurements (e.g. mean and s.e.m.) if each bin contained at least five measurements. We also only included bins if the measurements had movements with direction error relative to the target (defined as the difference in the angular direction of a saccade relative to the angular direction of the target displacement vector) of less than 45 deg (this was the great majority of data; e.g. Fig. 8 in Results).

For the re-analysis of the delayed, visually-guided and memory-guided saccade data of (Willeke et al. 2019), we used similar procedures to those described above. Since the sampling of target locations in these tasks was slightly different from that performed in the present experiments (for the visually-guided saccade task), we adjusted the binning windows accordingly, and we only included any measurement bins in which there were at least 7 saccades per bin. We also accepted as a minimum reaction time 100 ms instead of 70 ms, since we observed that reaction times in these “delayed” types of saccade tasks were generally longer than in the immediate visually-guided saccade task.

For the re-analysis of the human memory-guided saccade data of (Willeke et al. 2019), we again used similar procedures. Like in the monkey memory-guided saccade task, we considered a minimum reaction time of 100 ms. In reality, this was conservative, since the human reaction times were significantly longer, in general, than those of the monkeys in the same task (as described in Results and also in Willeke et al. 2019).

For the visually-guided saccade task, we also plotted saccade amplitude and direction error as a function of target eccentricity, using similar binning procedures to those described above.

In all analyses, we were interested in comparing upward and downward saccades, since eye-movement related structures like the superior colliculus (SC) represent them differently (Hafed and Chen 2016). We therefore divided trials according to whether the target location was downward (one group) or purely horizontal and upward (another group).

### Statistical analyses

Our purpose in the present study was to document patterns of rhesus macaque reaction time values as a function of visual-field location across a variety of well-established oculomotor tasks. We therefore present descriptive statistics in all figures, showing mean and s.e.m. measurements, as well as numbers of observations. All trends that we focus on are immediately visible in the mean and s.e.m. plots that we present.

### Data availability

All data presented in this paper are stored in institute computers and are available upon reasonable request.

## Results

### Monkeys exhibit increased saccadic reaction times for foveal target eccentricities

Our goal was to document the saccadic reaction times of rhesus macaque monkeys when target displacements are small. We were motivated by the fact that in humans, it is known that small-eccentricity target displacements are associated with increased saccadic reaction times (De Vries et al. 2016; Kalesnykas and Hallett 1996; 1994; Wyman and Steinman 1973). Figure 1 shows example eye position and velocity traces from one monkey (monkey M) when the target displacement was small (Fig. 1a) or when it was much larger (Fig. 1b). In both cases, the target displacement was to the right of central fixation, and we plotted horizontal eye position as well as radial eye velocity in the interval around target jump (labeled target onset in the figure). In both cases, a saccade was made to the target, which scaled appropriately in size with target eccentricity (also see Fig. 8 in a later section of Results). However, when the target eccentricity was small (Fig. 1a), the small saccades had significantly longer reaction times than the big saccades generated when the target eccentricity was large (Fig. 1b). This is illustrated in Fig. 1 with dashed vertical lines delineating the reaction times of the fastest small (Fig. 1a, blue dashed line) and large (Fig. 1b, red dashed line) saccades in the two shown data sets. As can be seen, there was a clear difference between fastest reaction times in the shown movements as a function of target eccentricity. Moreover, the overall distribution for the small saccades was shifted towards longer and more variable reaction times when compared to the bigger saccades. These examples demonstrate that rhesus macaque monkeys exhibit the same latency increase for small visually-guided saccades as human subjects (De Vries et al. 2016; Kalesnykas and Hallett 1996; 1994; Wyman and Steinman 1973).

**Figure 1.**
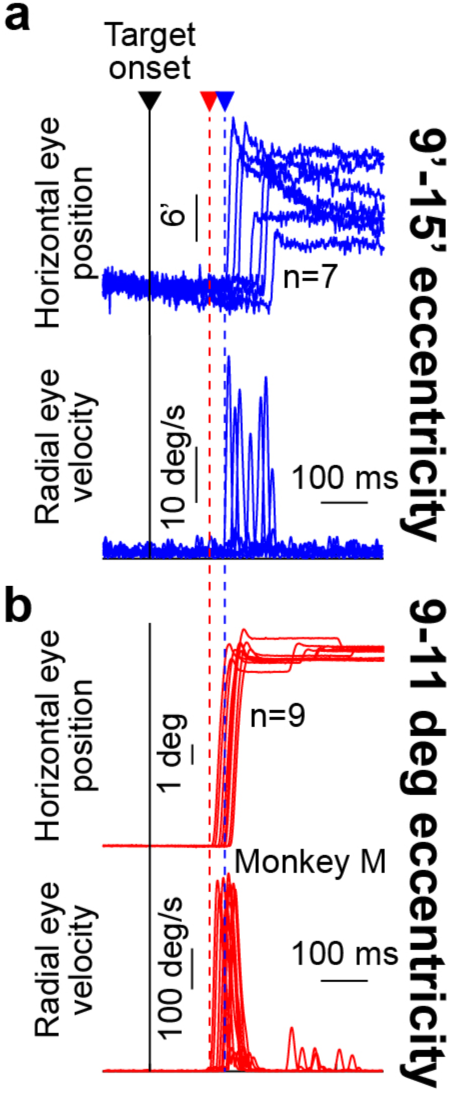
Example horizontal visually-guided saccades of different amplitudes in a rhesus macaque monkey. **(a)** Eye position (top) and velocity (bottom) measurements from 7 example trials in monkey M, for rightward target onsets at eccentricities between 9 and 15 min arc. Upward deflections in each eye position trace (top) mean rightward eye position displacements, and the position scale bar denotes 6 min arc. The vertical dashed blue line indicates the reaction time of the fastest saccade to occur in the shown set, to facilitate comparison to the data in **b**. **(b)** Similar analyses for 9 example movements from the same monkey, but now for target eccentricities between 9 and 11 deg, again for rightward target onsets. Note that the position scale bar (top) now indicates 1 deg. The vertical dashed red line indicates the reaction time of the fastest movement to occur in the shown set. Comparison of the blue and red traces (as well as the blue and red vertical dashed lines) reveals a clear increase in reaction times for the small saccades. Subsequent figures further characterize such an increase.

We summarized the above results across the entire population of measurements. In Fig. 2a, we plotted in the leftmost panel the mean (surrounded by s.e.m. boundaries) saccadic reaction time of monkey M as a function of target eccentricity. The smallest target eccentricities (<1 deg) were associated with long reaction times, reaching a mean of approximately 240 ms. Reaction time then dropped down to approximately 150 ms for eccentricities >1 deg. Larger eccentricities (approximately >5 deg) were associated with another increase in saccadic reaction times, albeit not as strong as that for the foveal target eccentricities. This strong increase for the foveal targets is more vivid in the middle panel of Fig. 2a, zooming in on only the central 1.5 deg of target eccentricities. Similarly, the rightmost panel of Fig. 2a plots the same data as in the leftmost panel but now on a logarithmic eccentricity scale, again demonstrating the longer saccadic reaction times associated with small target eccentricities. Similar results were obtained in monkey N, except that this monkey showed an even more dramatic increase in reaction times for foveal target eccentricities (from a minimum mean reaction time of <150 ms to a peak of approximately 300 ms). Therefore, across all target locations and eccentricities that we measured, there was a clear and consistent increase in saccadic reaction times of the two monkeys for foveal targets. There was another increase in reaction times, albeit weaker, for large target eccentricities, as also observed in human subjects (Kalesnykas and Hallett 1996; 1994).

**Figure 2.**
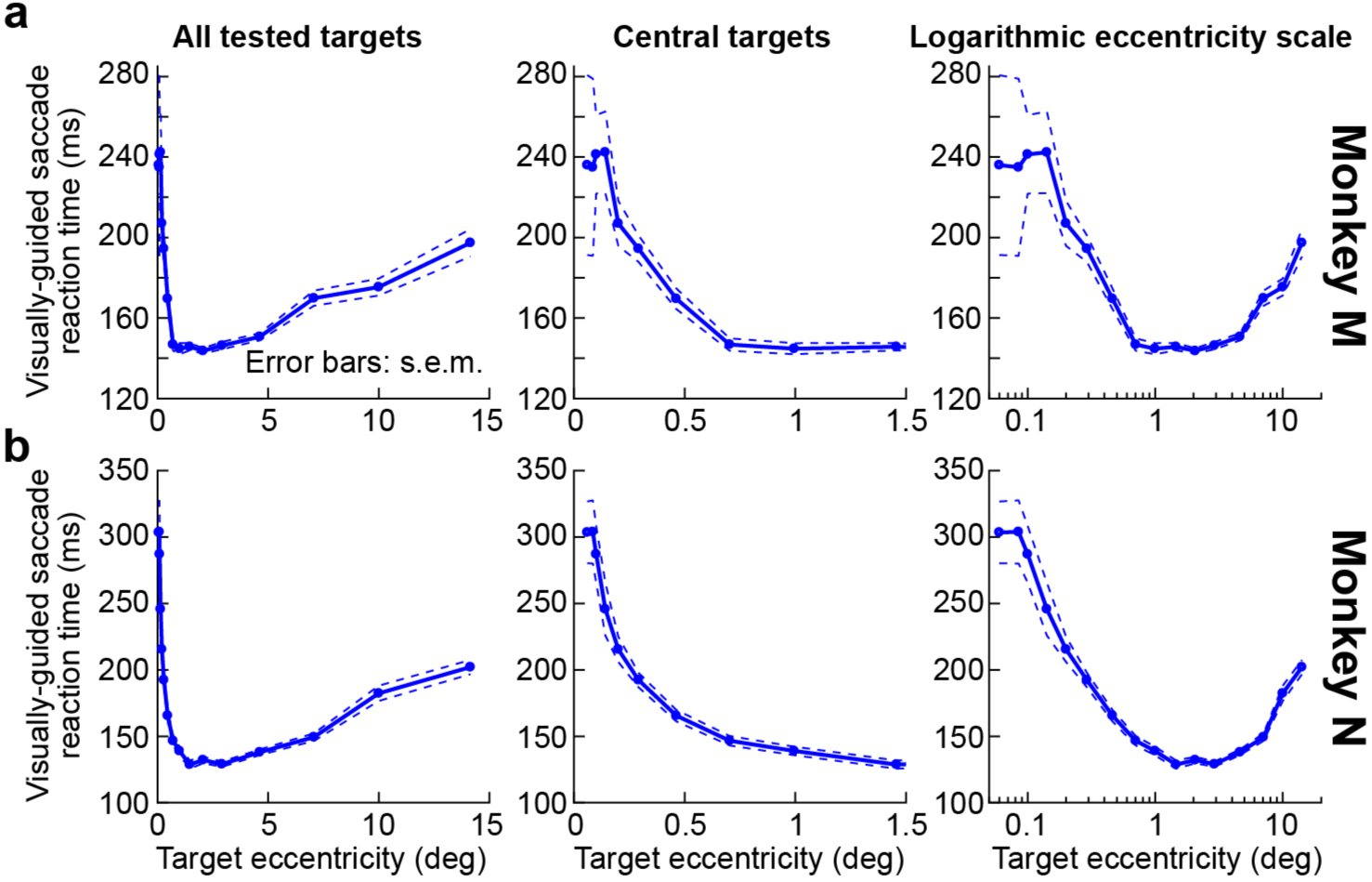
Longer visually-guided saccade reaction times for small target eccentricities in rhesus macaque monkeys. **(a)** In monkey M, we plotted visually-guided saccade reaction times as a function of target eccentricity (left). Reaction times for small-amplitude saccades were the longest. Zooming in to the central 1.5 deg (middle) revealed a monotonic decrease in reaction time with increasing amplitude within foveal target eccentricities. The rightmost plot shows the same data in the left and middle panels but using a logarithmic x-axis scale to clarify the strong increase in reaction time when small-amplitude saccades are triggered. Note that the reaction times also increased again for larger target eccentricities (e.g. > 5 deg). n=928 trials; error bars in each panel denote s.e.m. **(b)** Similar results from monkey N. There was an even stronger increase in reaction times for foveal target eccentricities. n=1246 trials.

We also inspected raw reaction time distributions to confirm that our method for accepting successful trials during the experiments did not artificially penalize specific ranges of reaction times. Specifically, our monkeys were rewarded based on the use of virtual, computer-controlled windows surrounding target location. If the eye position was not inside the virtual target window within 500 ms from target onset on a given trial (Methods), then the trial was aborted and the monkey was not rewarded. It is therefore conceivable (although unlikely; Methods) that we artificially truncated reaction time distributions at 500 ms, especially for target eccentricities showing increased reaction times (Fig. 2). However, this was not the case. For example, the top panels of Fig. 3a, b show the raw reaction time distributions of our two monkeys when foveal target eccentricities of 12-36 min arc were tested. The distributions were not truncated at 500 ms, suggesting that the monkeys were able to generate visually-guided saccades to these foveal targets within the prescribed “grace” period of 500 ms. Similarly, the bottom panels of Fig. 3a, b demonstrate that for another range of target eccentricities in which reaction times increased (Fig. 2), the increase was again not affected by the truncation at 500 ms forced by our grace period.

**Figure 3.**
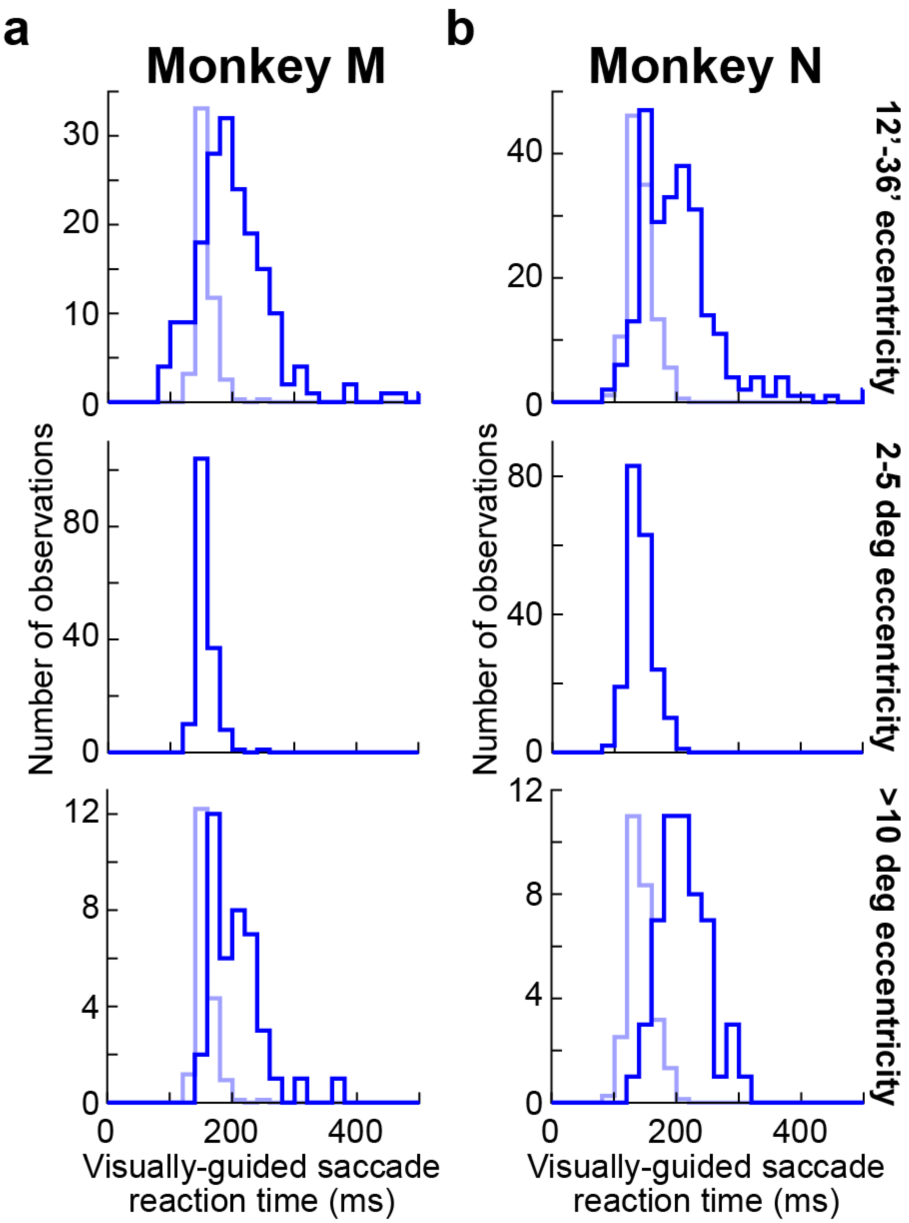
Raw distributions of visually-guided saccade reaction times for different representative target eccentricities. **(a)** In monkey M, we plotted the distributions of saccade reaction times for three ranges of eccentricities chosen based on the results of Fig. 2: the top panel shows data when target eccentricity was 12-36 min arc; the middle panel shows data when target eccentricity was 2-5 deg; and the bottom panel shows data when target eccentricity was >10 deg (and <20 deg; Methods and Fig. 2). The faint-colored curves in the top and bottom panels are copies (with arbitrary y-axes) of the curve in the middle panel to facilitate comparison of the distributions. Note how the distributions of the top and bottom panels demonstrate that the increased reaction times observed in Fig. 2 at these eccentricities were not due to a potential artifact caused by a cut-off grace period of 500 ms for responding with an eye movement (Methods). **(b)** Similar results from monkey N. In all panels, target directions (e.g. lower versus upper visual field locations) were equally distributed across all trials. Also note that numbers of observations are directly inferable from the histogram counts in each panel.

### Targets in the lower visual field are associated with increased reaction times even for foveal eccentricities

Because the SC exhibits a strong asymmetry between the representation of the upper and lower visual fields (Hafed and Chen 2016), with direct consequences on saccadic reaction times for large saccades, we next analyzed the reaction times of small visually-guided saccades to foveal targets in the upper and lower visual fields. Specifically, it was not known so far whether differences in saccadic reaction times between upper and lower visual field target locations also occur for very small eye movements. For the same data as in Fig. 2, we divided trials according to whether the target was in the lower visual field (Fig. 4a, c; red) or whether it was along the horizontal meridian or in the upper visual field (Fig. 4a, c; blue). Using the same formatting conventions as in Fig. 2, we found that there was, essentially, a global upward shift in the relationship between saccadic reaction time and target eccentricity for targets in the lower visual field. That is, the reaction time increase associated with lower visual field target locations also happened for tiny foveal eccentricities (middle panels in Fig. 4a, c). Moreover, this effect was not restricted to cardinal target/saccade directions. For example, in Fig. 4b, d, we plotted, for each monkey, the saccadic reaction time as a function of two-dimensional target location. We plotted target location bins on a log-polar axis (Hafed and Krauzlis 2012), in order to cover the large span of eccentricities tested, and we color-coded each binned target location (z-axis) with the mean saccadic reaction time for that location. In both monkeys, the same general dependence of saccadic reaction time on target eccentricity (Figs. 2, 3, 4a, c) occurred for all target directions. That is, foveal locations had the longest reaction times; there was a minimum of reaction times at intermediate eccentricities; and there was then a more modest increase in reaction times once again for larger eccentricities. Moreover, lower visual field locations (including foveal ones) were associated with the longest reaction times (also see Fig. 9 for a human replication of all of these observations).

**Figure 4.**
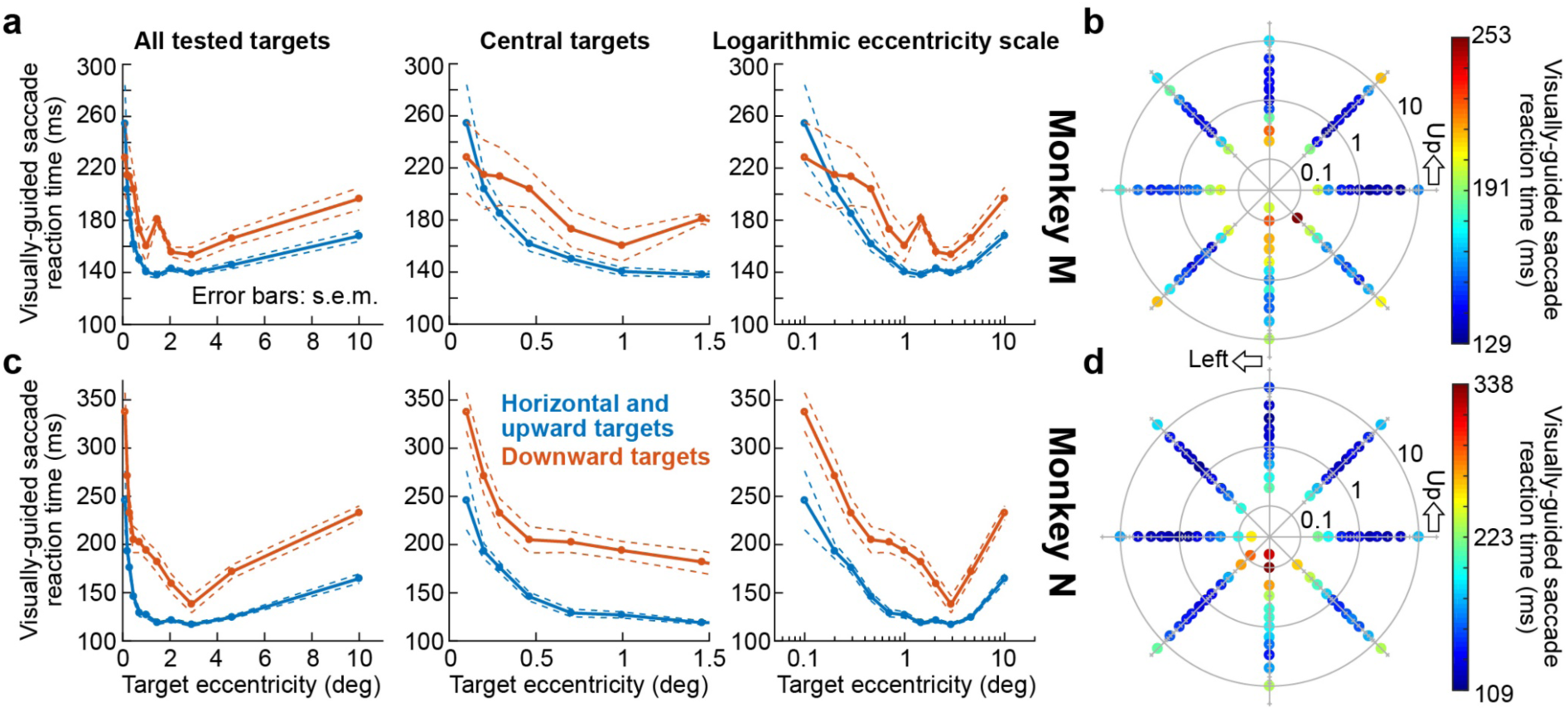
Dependence of monkey visually-guided saccade reaction times on upper versus lower visual field target location. **(a)** Similar analyses as in Fig. 2a for monkey M. However, here, we separated target locations as being either horizontal and upward relative to the line of sight (blue) or downward (red). Reaction times were longer for downward targets, consistent with (Hafed and Chen 2016). This also happened for small target eccentricities (middle panel of **a**). **(b)** Log-polar plot of the same data as in **a** demonstrating how intermediate eccentricities had the lowest reaction times in all directions, and how lower visual field locations were associated with longer reaction times than upper visual field locations at all eccentricities tested. **(c, d)** Same as **a**, **b** but for monkey N. Error bars denote s.e.m.

### Delayed, visually guided saccades show largely similar reaction time patterns to (immediate) visually-guided saccades

Previous experiments in humans have attempted to dissociate between visual and oculomotor (or other) sources of increased reaction times for small target eccentricities, and they attributed the increase to a difficulty in specifying the saccade metrics (Kalesnykas and Hallett 1996; Wyman and Steinman 1973). Similarly, in the rhesus macaque SC, while some visual bursts for foveal target onsets might show dependence on foveal eccentricity in their response latency (first-spike latency), this does not seem to be a general property of foveal SC neurons (Chen et al. 2019). Specifically, superficial SC neurons show decreases in first-spike latency of the visual response with increasing foveal eccentricity (consistent with our behavioral findings above), whereas deeper SC neurons show no such dependence of visual burst latency on foveal target eccentricity (Chen et al. 2019). Since it is the deeper SC neurons that show higher correlations between visual burst latency and saccadic reaction times (Chen and Hafed 2017; Marino et al. 2012), this might suggest that increased reaction times for small target eccentricities may not be intrinsically visual in nature (i.e. caused purely by visual-only mechanisms). Finally, other studies showed that the latencies of gaze shifts increase for the smallest gaze displacements, and not for the smallest target eccentricities (Goffart and Pelisson 1997; Zambarbieri et al. 1995), corroborating the view that saccadic initiation is dependent upon signals related to the impending movement rather than incoming signals from the retina. To test this, we ran our monkeys on a delayed saccade task. In this task, the target remained persistent while the fixation spot was visible. Only when the fixation spot was removed were the monkeys allowed to make the saccade (Methods).

We found a similar increase in reaction time in the delayed condition as in the immediate visually-guided saccade task for small target eccentricities. This happened even though the target was persistent, and the instruction to trigger the saccade was the offset of a fixation spot instead of the onset of the target. The task also had temporal expectation inherently built into it, since the longer the delay period was, the more likely it was that the “go” signal for the saccade was to come; there was also sufficient time with short delay periods to plan a saccade. Figure 5 plots reaction time data from this task in a format identical to that in Fig. 4 for both monkeys. The same general pattern of results was observed. Namely, small, foveal target eccentricities were associated with the longest reaction times, and lower visual field locations were also associated with long reaction times when compared to horizontal and upper visual field locations.

**Figure 5.**
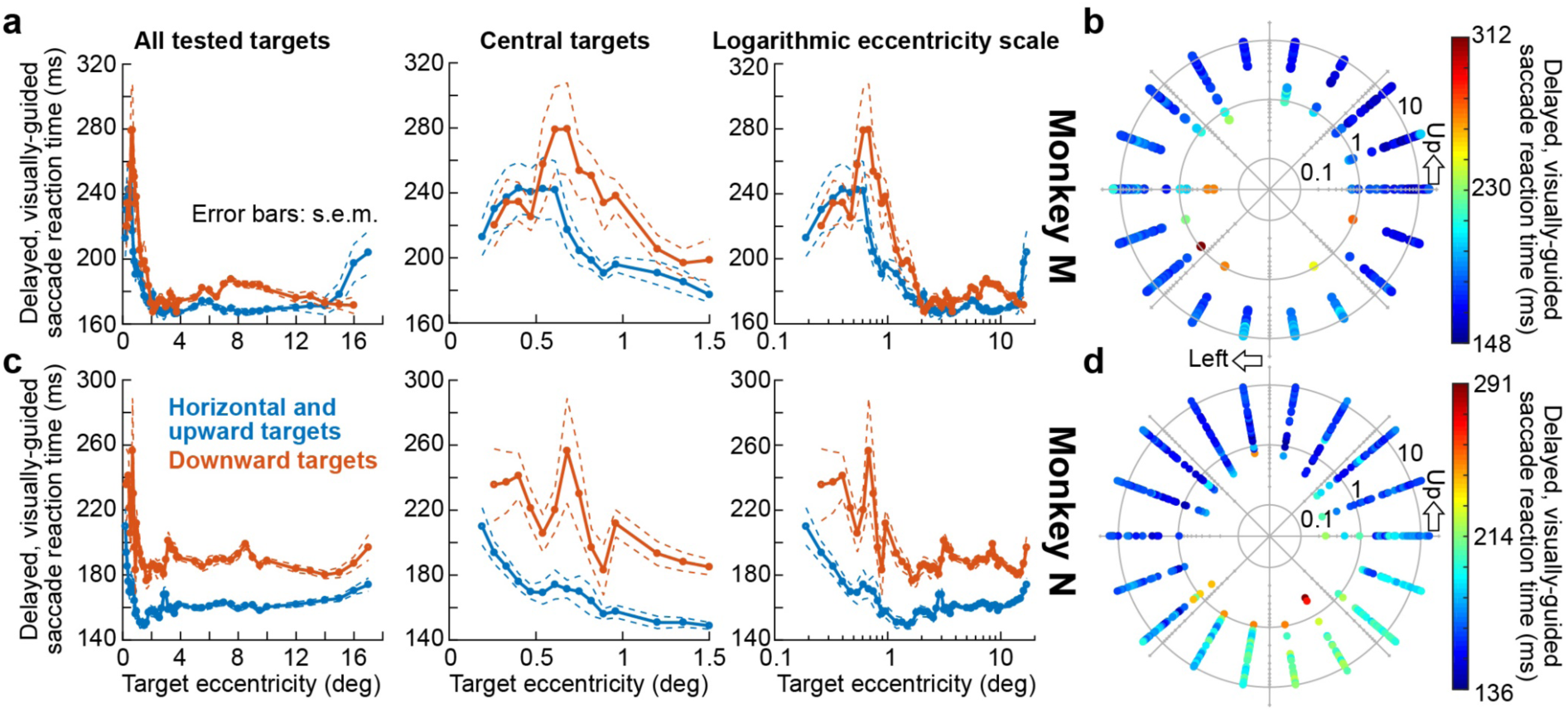
Increased reaction times for small delayed, visually-guided saccades in the rhesus macaque monkey. **(a)** Similar analyses to those in Fig. 4a for monkey M, but now during the delayed, visually-guided saccade task. As in the (immediate) visually-guided saccade task, small saccades had increased reaction times compared to larger saccades. Also like in the visually-guided saccade task, upper visual field targets were associated with faster reaction times than lower visual field targets. However, an increase in reaction times for large target eccentricities was less clear here than in the (immediate) visually-guided saccade task. Instead, downward targets of intermediate eccentricities (e.g. 5-12 deg; red reaction time curves in **a**) were associated with increased reaction times relative to eccentricity-matched upward targets. n=3265 trials; error bars denote s.e.m. **(b)** Log-polar plot showing the underlying data of **a** across space. (**c, d**) Similar results were obtained with monkey N. n=4277 trials.

An interesting difference that emerged in this condition relative to the (immediate) visually-guided saccade condition was the behavior of saccadic reaction times for large eccentricities (e.g. >10 deg). In this variant of the task, the increase in saccadic reaction times with increasing target eccentricities was less consistent than with the (immediate) visually-guided saccade task (Figs. 2-4). Instead, lower visual field targets of intermediate eccentricities (e.g. between ∼4 and 10 deg) exhibited a small increase in reaction time relative to larger target eccentricities (and upper visual field target locations). Thus, even though knowledge of target location and expectation to generate a saccade altered the detailed patterns of saccadic reaction times for extra-foveal target locations, the same increases in reaction times for foveal targets were evident in this task just like in the (immediate) visually-guided saccade task.

### Small memory-guided saccades are also associated with increased reaction times, despite the absence of a visual target

To further demonstrate the independence of small saccade reaction times in the rhesus macaque monkey from foveal visual responses (whether in SC or elsewhere), we also trained our monkeys to generate small memory-guided saccades (Willeke et al. 2019). In this case, the instruction to generate a saccade was the offset of a fixation spot displayed on an otherwise blank screen. The saccade itself was not directed to a visual stimulus, but instead to a remembered location (Willeke et al. 2019). We found similar increases in saccadic reaction times for foveal target eccentricities (Fig. 6; formatted identically to Figs. 4, 5). Interestingly, for foveal target eccentricities (middle panels of Fig. 6a, c), there was no clear difference in reaction times between locations in the upper and lower visual fields, unlike when there was a visual stimulus as the target for the saccade (Figs. 4, 5). Thus, even with memory-guided “microsaccades” (Willeke et al. 2019), there was an increase in saccadic reaction times, although the presence or absence of a visual target could alter the detailed properties of such an increase. It is also worth noting that the reaction time in this condition did not increase for larger eccentricities as in the (immediate) visually-guided saccade task. Instead, and as in the delayed, visually-guided saccade task, it was specifically the lower visual field saccades for intermediate eccentricities that seemed to increase.

**Figure 6.**
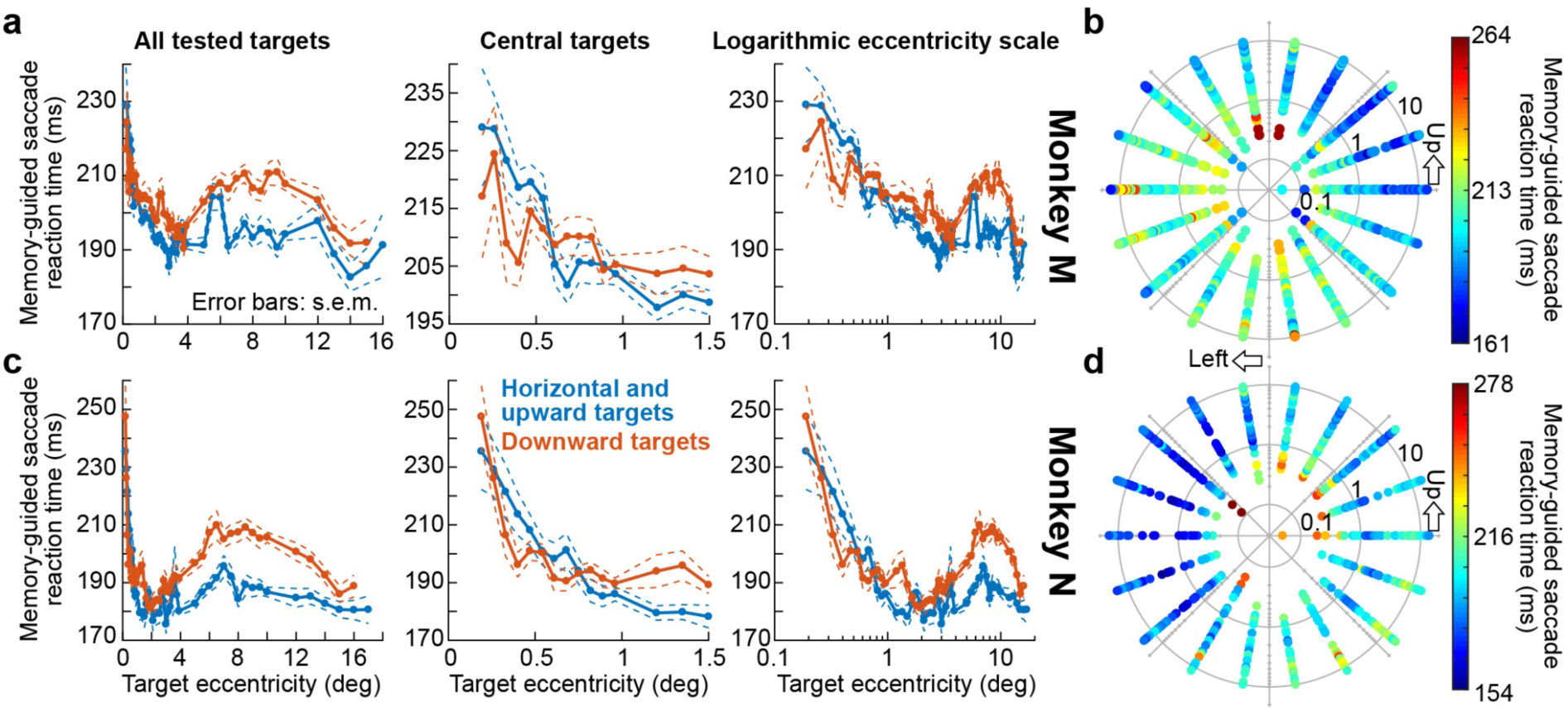
Increased reaction times for small memory-guided saccades in the rhesus macaque monkey. **(a, b)** Similar analyses to Fig. 4a, b for monkey M but during the memory-guided saccade task. Small memory-guided saccades were associated with increased reaction times (compared to larger saccades) despite the absence of a visual target for the saccades. Note that for memory-guided saccades, the increase in reaction time for larger target eccentricities that was obvious for (immediate) visually-guided saccades (e.g. Fig. 2) did not take place so clearly. Instead, downward saccades of intermediate sizes (e.g. 5-12 deg amplitudes; red curves) showed elevated reaction times compared to upward saccades. Also note that there was no difference in reaction time between upward and downward targets for small amplitudes (middle panel in **a**), unlike for visually-guided saccades (Figs. 2-5). n=6346 trials; error bars denote s.e.m. **(c, d)** Same results for monkey N. n=4245 trials.

### Small memory-guided saccades in humans show patterns to small memory-guided saccades in monkeys

Intrigued by the results documented in Fig. 6, we sought to test whether similar observations could also be made in human subjects. We had human subjects perform the same task as the monkeys (Willeke et al. 2019) and found very similar results (Fig. 7). Small memory-guided “microsaccades” (Willeke et al. 2019) were associated with the longest reaction times relative to all other eccentricities, just like in the monkeys. Therefore, in all tasks, small saccades were always associated with the longest latencies, regardless of whether the saccades were reflexive (Figs. 1-4), delayed (Fig. 5), or memory-guided (Figs. 6, 7).

**Figure 7.**
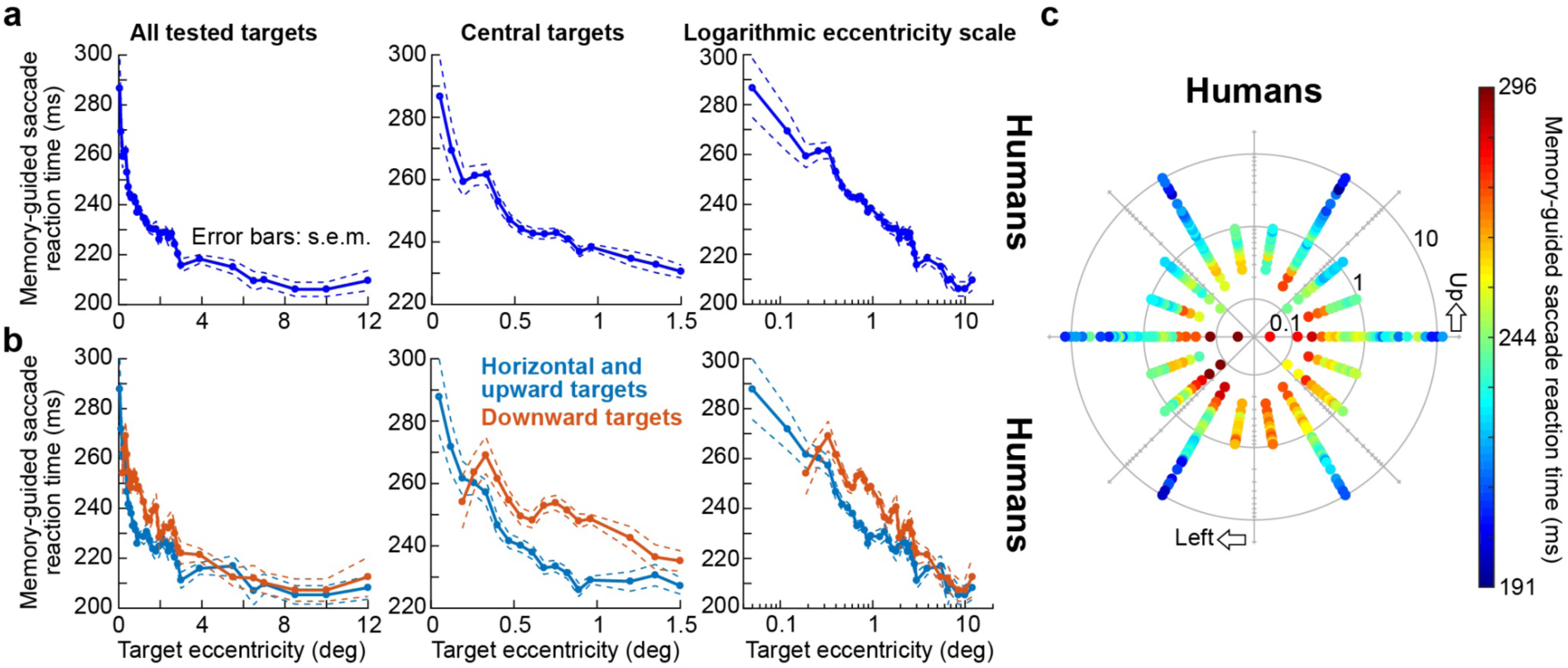
Memory-guided saccades in humans showed the same eccentricity-dependencies as monkey memory-guided saccades. **(a)** Similar analyses to Fig. 2 but for our human subjects’ data combined. The smallest saccades were associated with the longest reaction times, as in the monkeys (Fig. 6). n=13531 trials; error bars denote s.e.m. **(b)** The same data but now with upward and downward target locations separately, as in Fig. 6. Similar results to those in the monkeys were obtained. Note that for small amplitudes (middle panel), there was a clearer difference between upward and downward saccades than in the monkeys (Fig. 6). Thus, the data appeared more similar to those in visually-guided saccades (Fig. 4a, c; middle panels). **(c)** The same data but in a format similar to that of Fig. 6b, d, demonstrating that, in humans as well, small memory-guided “microsaccades” in all directions are associated with increased reaction times.

### Small visually-guided saccades show differences in spatial accuracy for upward and downward targets

Finally, because of the global changes in reaction times in the (immediate) visually-guided saccade task for different visual field locations (Fig. 4), we searched for other asymmetries in saccade parameters that also depended on foveal (or extra-foveal) target location. We found that saccade amplitude and direction differentially depended on visual field target location for foveal targets. Specifically, when we plotted saccade amplitude as a function of target eccentricity (Fig. 8a, c), we found that amplitude scaled nicely with target eccentricity even for foveal target locations, but there was larger overshoot in saccade amplitude for foveal targets in the lower visual field than in the upper visual field or along the horizontal meridian. On the other hand, when we plotted saccade direction error relative to target direction (Fig. 8b, d), we found that the overshooting lower visual field small saccades were more directionally accurate than the saccades to foveal targets in the upper visual field or along the horizontal meridian. Therefore, besides strong increases in reaction times for foveal target eccentricities (Figs. 1-4), small visually-guided saccades showed differential effects of amplitude versus directional accuracy as a function of target visual field location. These effects of Fig. 8 were not so clearly visible in the other variants of the task, such as the delayed, visually-guided saccade task or the memory-guided saccade task.

**Figure 8.**
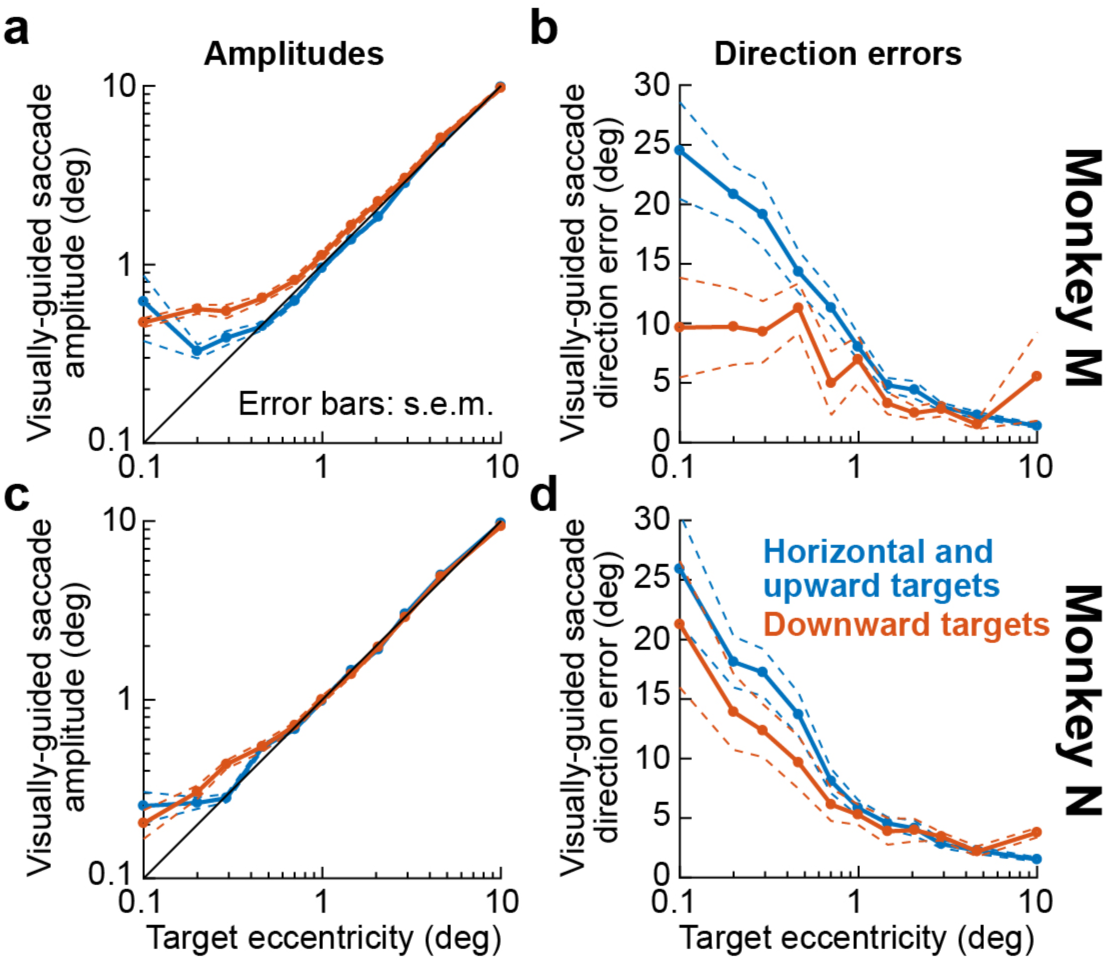
Small visually-guided saccades in the rhesus macaque monkey were more accurate in size for upward saccades but more accurate in direction for downward saccades. **(a)** Saccade amplitude as a function of target eccentricity in monkey M. We used logarithmic axes to display the data such that the effects with small eccentricities are more visible. Downward small saccades (red) overshot the target significantly more than upward and horizontal small saccades (blue). n=928 trials. **(b)** For the same data as in **a**, we plotted saccade direction error (relative to target direction) as a function of target eccentricity. Direction error was smaller for small downward saccades (red) than for small horizontal and upward saccades (blue). **(c, d)** Similar results for monkey N. n=1246 trials. In all panels, error bars denote s.e.m.

## Discussion

We investigated spatial and temporal aspects of triggering small saccades in the rhesus macaque monkey and compared our results to those obtained in human subjects both from the existing literature and through our own additional measurements. We specifically found that, in the monkey, small saccades require more time than larger saccades to be triggered. This observation is true whether the small saccades are visually-driven, delayed (but still visually-driven), or memory-guided. We also found a strong asymmetry in the reaction times of small upward versus downward visually-guided saccades, similar to larger saccade results (Hafed and Chen 2016; Schlykowa et al. 1996; Zhou and King 2002). For larger saccades, there was a gradual increase in reaction times with increasing target eccentricities, but primarily in the (immediate) visually-guided saccade task and not in the delayed, visually-guided saccade task or the memory-guided saccade task.

Our results are important to document in the oculomotor system literature because there has been no systematic attempt to investigate small-saccade triggering in the rhesus macaque monkey. In humans, it is well-known that small-saccade reaction times are long (Kalesnykas and Hallett 1996; 1994; Wyman and Steinman 1973), although the mechanistic reasons for this phenomenon seem to be still not fully understood. In the monkey, there have basically been only casual inferences about small-saccade reaction times in macaques (Boch et al. 1984; Skinner et al. 2019; Willeke et al. 2019), but either with tasks that were not explicitly designed for such analyses or without sufficient sampling of small target eccentricities. Therefore, our results provide an important reference catalog of small-saccade triggering in macaque monkeys. This is especially important nowadays to guide investigations of neural mechanisms associated with foveal visual and motor processing (Chen et al. 2019; Guerrasio et al. 2010; Willeke et al. 2019).

Our results are additionally important because other primate species, like marmoset monkeys (Sedaghat-Nejad et al. 2019), are starting to be used for studying the oculomotor system. Even though rhesus macaques remain the reference model because of the history of their usage, it is important to identify the diversity of oculomotor capabilities across various non-human primates in order to bridge the gap between the knowledge acquired from genetic, neuroanatomical, neurophysiological, and behavioral studies.

One interesting aspect of our results is the observation that the reaction times of small upward saccades are shorter than those of small downward saccades (e.g. Fig. 4). This was in addition to the observation of increased reaction times in general for small saccades (Figs. 1-3). Therefore, not only are foveal targets associated with long saccadic reaction times (Figs. 1-3), but longer times are particularly prominent with foveal targets located in the lower visual field (Fig. 4). Interestingly, in one of their control conditions, Wyman and Steinman (1973) required a small downward saccade, which exhibited prolonged reaction times in humans as well, although this aspect of the data was not explicitly mentioned in that study (see Fig. 9 below for further evidence). These effects are consistent with the asymmetric representations of upper and lower visual fields in the SC, in such a manner that can directly affect saccadic reaction times and landing errors (Hafed and Chen 2016). These effects are also consistent with our theory of gaze direction as an equilibrium insofar as an eye movement is not initiated as long as the visuo-oculomotor system is within a state of balance among opposing commands (Goffart 2019; Goffart et al. 2018; Goffart et al. 2012; Krauzlis et al. 2017).

**Figure 9.**
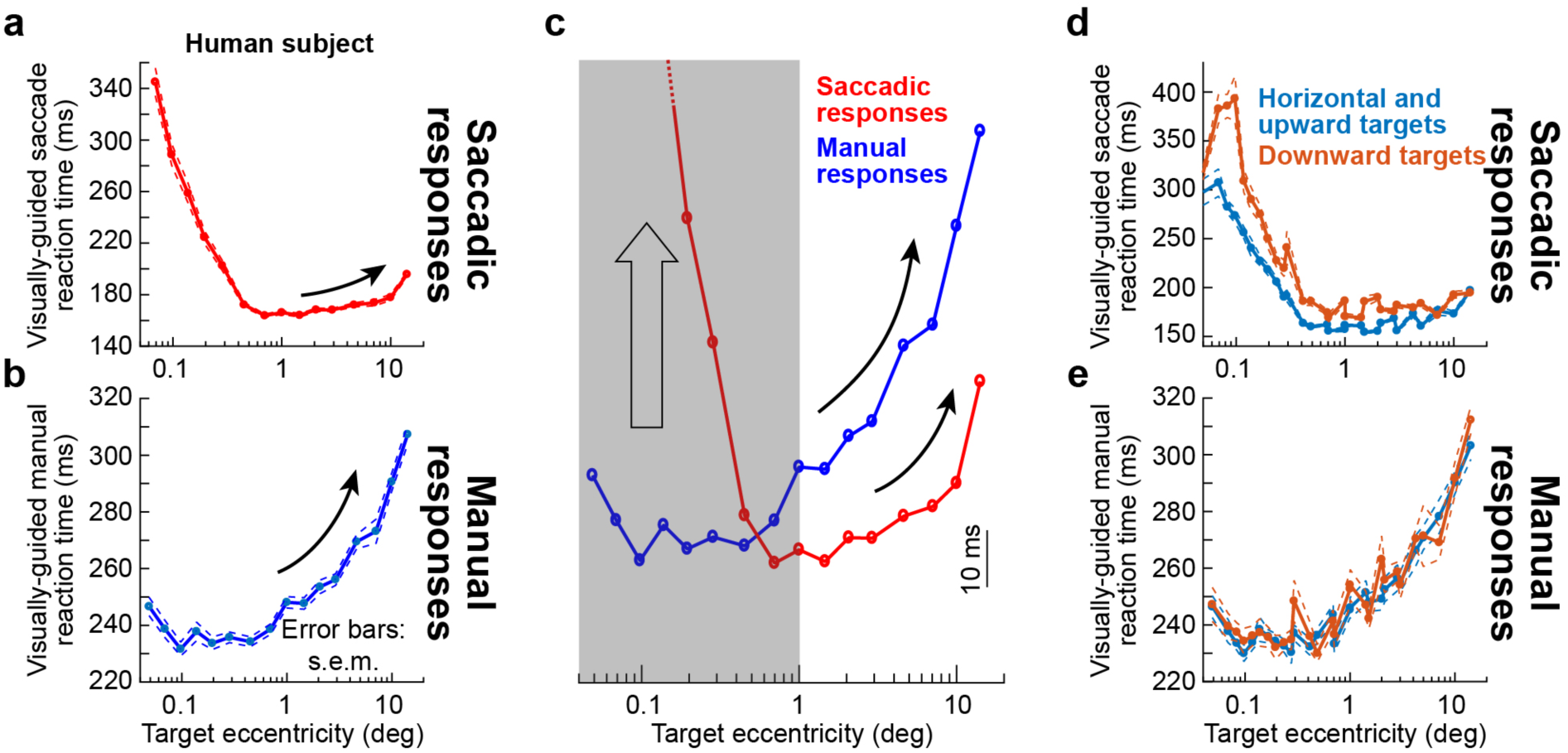
Both saccadic and manual reaction times increased for large target eccentricities in a human subject; only saccadic, but not manual, reaction times increased for small saccades as well as downward target locations. **(a)** We replicated well-known human findings (Kalesnykas and Hallett 1996; 1994; Wyman and Steinman 1973) in an expert subject (author ZH). The subject performed the same task as in Fig. 2 for our two monkeys. Note how small saccades had very long reaction times. Large saccades also showed a modest reaction time increase (curved black arrow). **(b)** When the same subject maintained fixation and detected target onset with a speeded manual reaction (button press), reaction times for foveal target eccentricities did not exhibit a strong increase. However, the large target eccentricities had the same manual reaction time increase as in **a** (curved black arrow). Note that the y-axis scale is different from **a**, visually masking the similarity between the two panels (but see **c** for a direct comparison). **(c)** To better show the similarity of reaction time increases for large target eccentricities in saccadic and manual responses, we plotted the same data from **a**, **b** using the same y-axis scaling (see vertical scale bar) but arbitrary y-axis placement to visually align the two curves. Note how both tasks showed an increase in reaction times for large target eccentricities (black curved arrows), but only saccadic responses showed strong increases for small target eccentricities (gray rectangle). **(d)** For the same data as in **a**, we separated upper and lower visual field target locations, as in Fig. 4. The human subject showed the same effects as our monkeys (Fig. 4). **(e)** However, manual reaction times did not show visual field asymmetries. Error bars denote s.e.m.

We also noticed interesting contrasts between the reaction times of small and large saccades. For example, immediate visually-guided saccades showed a marked increase in reaction times for large saccades (e.g. Fig. 2), but this effect was not as strong as that for small saccades. It has been reported in humans that large saccades also become associated with increased reaction times (Kalesnykas and Hallett 1996; 1994). However, are the causes similar to the causes of increased reaction times of small saccades? This issue remains unaddressed. Based on our data, we think that the two increases are driven by quite different underlying causes. Specifically, in the delayed and memory-guided saccade tasks, the increase in reaction times for large saccades was much less obvious than in the (immediate) visually-guided saccade task, even though small saccades showed strong increases in all three tasks. Therefore, increased reaction times for large saccades are driven by different mechanisms from those affecting small-saccade reaction times.

In our view, the absence of increase in reaction times for large saccades made in the memory-guided task suggests that the increase in the visually-guided task might critically depend on the perceptual detectability of the appearing target. Specifically, we used a small spot as the target in all of our experiments, even for eccentricities of 10 or 15 deg. This means that, at these eccentricities, the small spot would be harder to detect than in the foveal or parafoveal regions. Therefore, even without saccadic responses, decreased perceptual detectability at far eccentricities could delay reaction times. We explicitly tested this hypothesis by performing additional experiments with one of us (ZH) being the experienced subject (similar to our two experienced monkey subjects). Specifically, in Fig. 9a, we replicated the findings of human saccadic reaction times as a function of target eccentricity (Kalesnykas and Hallett 1996; 1994). Note how, in addition to the dramatic increase for small saccades, saccadic reaction time increased with increasing target eccentricity for extra-foveal targets (curved black arrow in the figure; similar to our monkeys in Fig. 2). Critically, for Fig. 9b, we ran the same subject on a perceptual detection task, in which no saccade was required at all. Instead, the subject had to press a button as soon as the target appeared in the periphery (Methods), and we confirmed that no microsaccades occurred between target onset and button press. Two notable observations are clear from the data. First, there was no increase in reaction times for foveal target eccentricities, suggesting that the increased reaction times of small saccades are specific to the fact that the motor behavior was to generate small saccades. Second, the same increase in reaction times for larger target eccentricities as in Fig. 9a was still evident (curved black arrow). This latter observation is much more obvious when the two curves are superimposed together in Fig. 9c using the same y-axis scaling (but with arbitrary y-axis positioning of the curves due to the different absolute values of saccadic and manual reaction times). As can be seen, both tasks were associated with increased reaction times for peripheral targets, but only the saccadic task was associated with increased reaction times for foveal targets. Therefore, the increases in Fig. 2 and Fig. 9a for large eccentricities were not specific to saccade generation. Interestingly, the human subject also showed the dependence of saccadic reaction times on upper versus lower visual field target locations (Fig. 9d) that we observed in our monkeys (Fig. 4), but this effect was again specific only to saccade generation (Fig. 9e). This is consistent with our interpretation above, and elsewhere, that asymmetries in upper and lower visual field representations in the SC have direct functional consequences on the oculomotor system (Hafed and Chen 2016).

If perceptual detectability can affect reaction times of large target eccentricities (Fig. 9), we think that equilibrium states in the oculomotor system explain the long reaction times of small saccades. Specifically, evidence from pharmacological inactivation experiments in either the SC or the caudal fastigial nucleus suggests that gaze direction is an equilibrium, and that an eye movement (saccadic or slow) is not initiated as long as the visuo-oculomotor system is within a state of balance where opposing commands counterbalance each other (Goffart 2019; Goffart et al. 2018; Goffart et al. 2012; Krauzlis et al. 2017). Thus, the longer reaction times of small saccades are the signature of enhanced times to create a symmetry-breaking condition, which puts the downstream premotor neurons into a push-pull regime that is responsible for the firing rates of motor neurons innervating the agonist and antagonist extraocular muscles (reciprocal innervation).

Consistent with the large amount of foveal magnification in the SC (Chen et al. 2019), this equilibrium idea is the extension of the population coding of saccade metrics to the specification of gaze direction during fixation. On the basis of recordings made in the SC, Sparks and colleagues indeed proposed that “precise saccadic movements are not produced by the discharge of a small population of finely tuned neurons but result from the weighted sum of the simultaneous movement tendencies produced by the activity of a large population of less finely tuned neurons” (Sparks et al. 1976). If the deep SC is a continuum representing different movement tendencies, then we are led to consider the absence of movement during fixation as the simple result of simultaneous movement tendencies counterbalancing each other. Thus, the foveal magnification in the SC contributes not only to ensure precise microsaccades (Chen et al. 2019; Guerrasio et al. 2010; Ko et al. 2010; Tian et al. 2018), but it does also imply that a large population of neurons is active at any one moment in time to represent the current fixated goal (Goffart et al. 2012; Hafed et al. 2008; Hafed and Krauzlis 2008). Therefore, a substantial shift is needed in such a large population to create sufficient imbalance that will excite one of the four populations of premotor neurons responsible for generating leftward, rightward, upward, and downward saccades.

If a balance among multiple gaze commands is what maintains gaze stability and increases reaction times for small saccades, then one might wonder how an imbalance may be generated at all during fixation in the first place. In other words, why is reaction time not infinite once balance among competing gaze shift commands is achieved? One possible explanation is behavioral and invokes the slow fixational eye movements that often happen in between saccades. These ocular drifts change the retinotopic location of the fixated object and thus create the imbalance needed to activate the colliculoreticular streams innervating the eye muscles. In fact, we recently found that the generation of tiny microsaccades during fixation is highly consistently associated with small, instantaneous retinotopic gaze position errors, even in the presence of peripheral “attention-capturing” cues (Tian et al. 2018; 2016). Another explanation is physiological and lies upon the fluctuations of the activity of neurons, which, from the foveal ganglion cells to the saccade-related premotor burst neurons, exhibit sustained firing rates whenever gaze is held stable. Among this immense number of neurons, we find not only long-lead burst neurons, but also neurons in the caudal fastigial nucleus (Kleine et al. 2003; Sun et al. 2016). After unilateral inactivation of this nucleus, fixational saccades are not only dysmetric (Guerrasio et al. 2010), but the direction of gaze during fixation and pursuit is also always deviated towards the lesioned side (Bourrelly et al. 2018; Goffart et al. 2004; Guerrasio et al. 2010). The fact that the head can also be deviated following a unilateral SC or fastigial lesion indicates that the balancing of activity is a process which is not restricted to the determination of eye gaze direction (Goffart et al. 2018); head direction is also an equilibrium. Thus, the proposal has been made that gaze and head movements, instead of reducing putative signals encoding the angular distance between gaze (or head) and target locations in physical space, are separate processes which consist of restoring symmetries (Goffart 2019).

An interesting additional consequence of the oculomotor balance idea is that we can predict express, rather than slow, reaction times for small saccades under the right conditions related to instantaneous gaze position error. Indeed, so-called “express microsaccades” can happen when peripheral stimulus onsets momentarily bias a state equivalent to “unstable equilibrium” out of equilibrium (Tian et al. 2018).

Finally, it would be interesting that future experiments investigate the neural mechanisms for learning to fixate visual objects under different visual and oculomotor conditions, and in which various amounts of asymmetry would be incorporated. For example, if target shape is changing, whether for the currently fixated position or for next target positions, what are the consequences on the oculomotor balance? How does this change alter the SC and caudal fastigial nucleus population activity? It will also be important to characterize the neural mechanisms underlying the coordination between saccades and slow eye movements (including the ocular drifts), with or without a concurrent head movement.

## Acknowledgements

ZMH was funded by the Werner Reichardt Centre for Integrative Neuroscience (CIN). The CIN is an Excellence Cluster (EXC307) funded by the Deutsche Forschungsgemeinschaft (DFG). ZMH was also supported by the Hertie Institute for Clinical Brain Research at Tuebingen University, and DFG-funded Research Unit (FOR1847; project: HA6749/2-1). LG was supported by CNRS. We thank Aya Tarek, Antimo Buonocore, and Konstantin Willeke for help with data collection.

## Author contributions

ZMH collected the data. ZMH and LG analyzed the data and wrote the manuscript.

